# Evaluation of colorectal cancer subtypes and cell lines using deep learning

**DOI:** 10.1101/464743

**Authors:** Jonathan Ronen, Sikander Hayat, Altuna Akalin

## Abstract

Colorectal cancer (CRC) is a common cancer with a high mortality rate and a rising incidence rate in the developed world. The disease shows variable drug response and outcome. Molecular profiling techniques have been used to better understand the variability between tumours as well as cancer models such as cell lines. Drug discovery programs use cell lines as a proxy for human cancers to characterize their molecular makeup and drug response, identify relevant indications and discover biomarkers. In order to maximize the translatability and the clinical relevance of in vitro studies, selection of optimal cancer models is imperative. We have developed a deep learning based method to measure the similarity between CRC tumors and other tumors or disease models such as cancer cell lines. Our method efficiently leverages multi-omics data sets containing copy number alterations, gene expression and point mutations, and learns latent factors that describe the data in lower dimension. These latent factors represent the patterns across gene expression, copy number, and mutational profiles which are clinically relevant and explain the variability of molecular profiles across tumours and cell lines. Using these, we propose a refined colorectal cancer sample classification and provide best-matching cell lines in terms of multi-omics for the different subtypes. These findings are relevant for patient stratification and selection of cell lines for early stage drug discovery pipelines, biomarker discovery, and target identification.

## Introduction

Colorectal cancer (CRC) accounts for 10% of cancer related deaths^1^, amounting to approximately 1.4 million new colorectal cases and 693,900 deaths reported in 2012^2^. CRC is not a homogeneous disease and can be classified into different subtypes based on molecular and morphological alterations^3^. The disease occurs when normal epithelium cells transform to cancer cells as a result of acquired genetic and epigenetic alterations. Mutations in the WNT signaling pathway are thought to initiate the transformation to cancer^4,5^. This is followed by deregulation of other signaling pathways such as MAPK, TGF-*β*, and PI3K–AKT via acquired mutations^5,6^. Since the original description of the molecular pathogenesis of CRC, multiple additional pathways, mutations, and epigenetic changes have been implicated in the formation of CRC^7^. Based on integrative analysis of genomic aberrations observed in the TCGA samples, CRC tumors can be divided into hyper-mutated (≈ 16%) and non hyper-mutated (≈ 84%) cancers. The hyper-mutated cancers have microsatellite instability (MSI), resulting from defective mismatch repair or DNA polymerase proof-reading mutations^3^. The non-hypermutated, microsatellite stable (MSS) cancers, are characterized by chromosomal instability (CIN), with high occurance of DNA copy number alterations, and mutations in the APC, TP53, KRAS and BRAF genes^3^. In addition, most CRC tumors have aberrantly methylated genes, a subset of which could play a functional role in CRC^7^. A further subset of CRC tumors display a CpG island methylator phenotype (CIMP), where some tumor suppressor genes could be inactivated epigenetically^8^. The diversity of molecular disease mechanisms creates distinct molecular subtypes in CRC with different survival rates and responses to therapy. Different molecular subtyping schemes based on gene expression profiles were recently studied and summarized as the consensus molecular subtypes (CMS)^9^, designating four main CRC subtypes with distinguishing features. The CMS1 subtype is defined by hypermutation, microsatellite instability and strong immune activation; CMS2 is defined by chromosomal instability (CIN), WNT and MYC signaling activation; CMS3 is defined by metabolic dysregulation; and CMS4 is defined by growth factor *β* activation, stromal invasion, and angiogenesis^9^. However, approximately 13% of the tumors do not belong to a consensus subtype, as they have mixed gene expression signatures. They may represent distinct tumor types, or samples with intra-tumor heterogeneity. While the CMS classification is based on gene expression, follow-up analysis of the tumors revealed distinct copy number profiles, mutation frequencies, and methylation profiles^9^ (Figure S3), indicating that other omics types contain important information.

We propose a multi-omics method that can incorporate gene expression, copy number, and mutation data to identify CRC subtypes, with implications for patient stratification. In addition, our method is able to match cell lines to each subtype. In the future, it can also be used to assign best-matching xenografts or organoid models to the study of each subtype. This would be highly useful, as these have been shown to predict drug response in patients^10^. By leveraging multi-omics datasets, CRC samples that can not be associated to a CMS subtype are also assigned to an appropriate subtype. In addition, multi-omics signatures, incorporating gene expression, point mutations and copy number alterations, are a direct output of the method, and will not need to be generated post-hoc, e.g. by examining mutation rates in groups defined by gene expression profiles, as in the CMS. With these goals in mind, we used deep-learning on multi-omics data in order to refine CRC subtypes in an unsupervised manner, and match them to cell lines.

Latent factor analysis is an *unsupervised learning* technique. Genomic assays are high dimensional (tens of thousands of genes), and high dimensional spaces are challenging to analyze. This problem is further exacerbated by the introduction of multiple assays from different omics platforms. Latent factor analysis seeks to learn (infer) a lower dimension representation of the data, which preserves the important structure / patterns therein. This is sometimes referred to as dimensionality reduction. By describing the data using a handful of *latent factors*, rather than tens of thousands of genes, down-stream analysis, such as distance calculations and clustering, are simplified^11^. It is further desirable that latent factors be interpretable in the biological context, i.e. that the patterns summarized by a latent factor implicate cellular processes. The workhorse for latent factor analysis for multi-omics, as well as general data analysis, has been matrix factorization^12^. Latent factor analysis for multi-omics data typically includes concatenating the different omics data to a single matrix and applying a well-known matrix factorization algorithm, sometimes with weighting of the individual data sets. Multifactor analysis (MFA)^13^ and iCluster+^14^ are examples of such methods. Some such algorithms, such as MFA, impose orthogonality of factors, i.e. that the factors explain disjoint underlying processes, as in PCA. Orthogonality might be conceptually appealing, but is not a biological necessity. Orthogonal latent factors may be the best for statistical reconstruction of a dataset, and still be biologically difficult to interpret. On the other end of the spectrum, there are deep learning based methods which work as dimensionality reduction techniques that can deal with non-linearity and can generalize well on different problems. In addition, they can be sparse, i.e. each latent factor only depends on a few of the input genes, and each sample is described by only a handful of latent factors. Sparsity in the relationship between input genes and latent factors simplifies the task of biological interpretation of the model, that is, implicating known biological processes underlying the latent factors; sparsity in the relationship between latent factors and samples simplifies down-stream analysis such as clustering. The disentangled variational autoencoders (*β*-VAE)^15^ have the previously mentioned characteristics and as a deep learning framework, they have flexible architectures that can work with most problem sets. A similar method has been shown to be able to stratify cancers by their tissue type based on gene expression profiles^16^, and other autoencoder architectures have been used to integrate multi-modal data in robotics^17^, as well as protein function prediction^18^.

We use a multi-modal, stacked *β*-VAE to extract latent factors which are important for defining subtypes and predicting patient survival. We call the method, “Multi-omics Autoencoder Integration”, or, maui. We compare maui’s performance to state-of-the-art multi-omics analysis methods, and show that maui outperforms the matrix factorization methods. In addition, we map CRC cell lines to the latent factor space defined by maui. This allows us to use patterns recognized by maui as significant in CRC to rank in-vitro models’ suitability for studying tumors. We then assign suitable cancer cell lines for drug target studies aimed at different colorectal cancer subtypes.

## Results

### Refining CRC subtypes using multi-omics data

The CRC cohort in the TCGA data-set (n=519, See Materials and Methods) has been extensively studied, and a state-of-the-art subtyping scheme exists in the “Consensus Molecular Subtypes” for colorectal cancer, or CMS^9^ (Table 1). In order to validate that the latent factors learned by maui capture patterns relevant to cancer biology, we tested how well the latent factors recapitulate the known subtypes.

**Table 1.** Summary of TCGA tumors’ CMS labels.

| CMS label | Description | # Samples |
| --- | --- | --- |
| CMS1 | MSI Immune | 61 |
| CMS2 | Canonical | 175 |
| CMS3 | Metabolic | 60 |
| CMS4 | Mesenchymal | 123 |
| **Total with CMS label** |  | **419** |
| Without CMS label |  | 100 |
| **Total** |  | **519** |

We extracted latent factors from gene expression, point mutations, and copy number alterations using maui, as well as other published methods for multi-omics integration by dimensionality reduction: MOFA^19^, and iCluster+^14^. In order to quantify the relationship between latent factor representations and the CMS subtype to compare the methods, we used Support Vector Machines (SVM, See Methods) to assign a CMS label to each tumor based on their latent factor representation. We then computed ‘Receiver Operating Characteristics’ (ROC), and compute the area under the curve (See Methods). The area under the ROC (auROC) is a measure of classification accuracy, with a score of 0.5 being expected from random guessing, and 1.0 being perfect. All methods produce latent factors with some correlation to the CMS labels (Figure 1A). Using SVM, maui (auROC 0.98) marginally outperforms MOFA (auROC 0.96) and both maui and MOFA dramatically out-perform iCluster+ (auROC 0.73) (Figure 1B, Figure S1).

**Figure 1.**
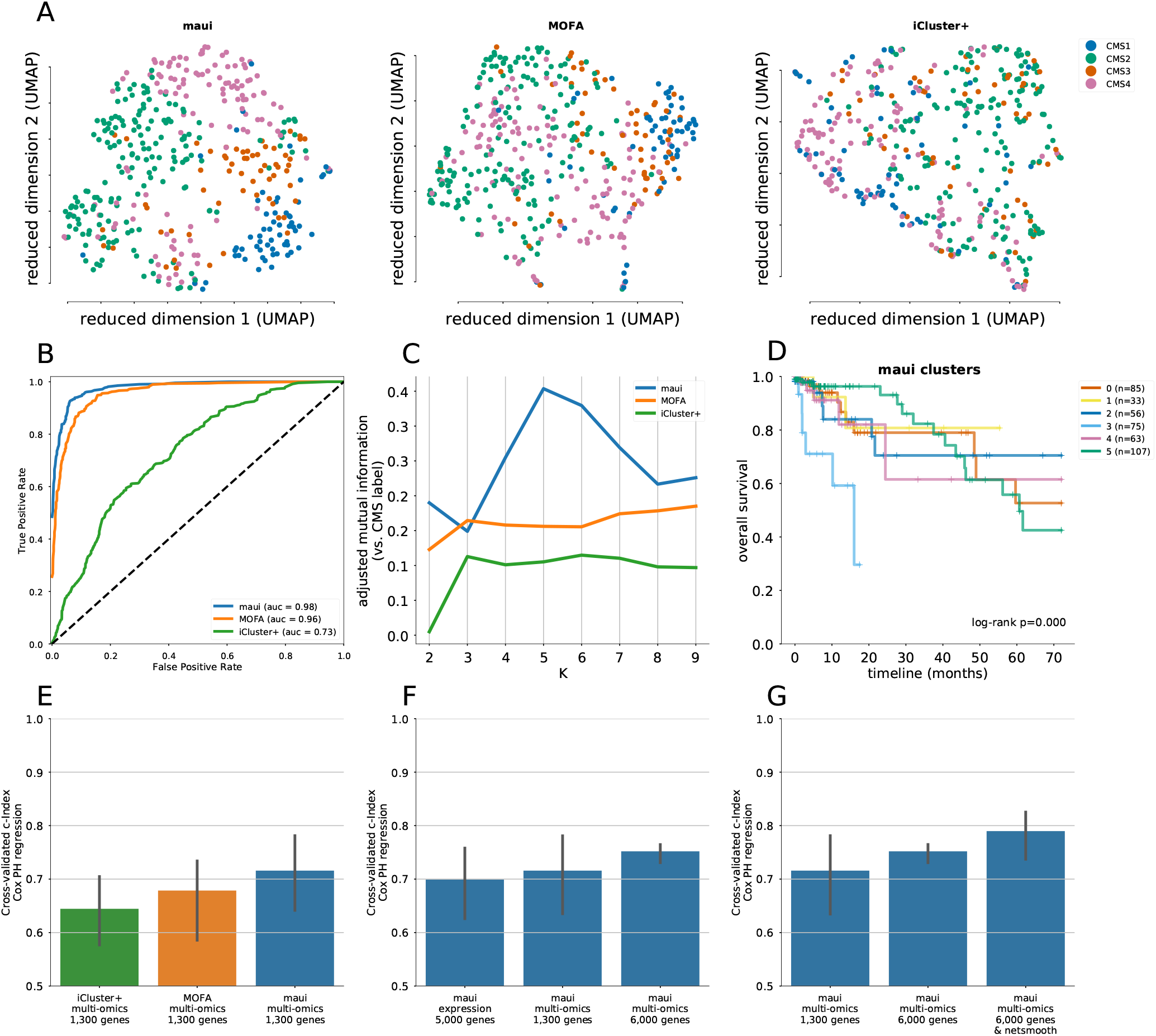
maui, MOFA, iCluster+, and the CMS labels. **A)** UMAP^24^ reduced dimensions from latent factors inferred by maui, MOFA, and iCluster+. Each dot represents a tumor, colored by their CMS label. **B)** ROC’s for regularized SVM’s predicting the CMS label from latent factors (out-of-sample, 10-fold CV). Mean ROC shown (See Methods) **C)** The Adjusted Mutual Information (AMI, See Methods) of clusters obtained from latent factors inferred by maui, MOFA, and iCluster+, using k-means clustering with K ranging from 2 to 9. **D)** Kaplan-Meier estimates and the log-rank statistic for differential survival of different clusters. The reported P value is from a multivariate log-rank test, under a null hypothesis that all groups have the same survival function. Clusters 3 and 5 represent a novel splitting of a previously defined subtype, CMS2. **E)** Harrell’s c-Index for Cox regressions of iCluster+, MOFA, and maui shows maui is more predictive of patient survival than other methods. **F)** Harrell’s c-Index comparing different maui flavors shows that maui benefits from multi-omics data, as well as from more input genes. **G)** Harrell’s c-Index shows network smoothing of mutations improves survival prediction using maui.

maui has an unfair advantage over MOFA in the previous analysis, as we ran it with 80 latent factors, whereas MOFA was only run with 20, and regularized supervised learning algorithms may benefit from a larger number of input features (here, the latent factors). In order to assess which of the methods is best able to capture the CMS labels, without regard to the number of latent factors, we repeated the previous exercise—predicting the CMS from the latent factors—using an *unsupervised learning* algorithm. We clustered the samples with a well-defined CMS^i^ using k-means clustering on the latent factors (See Methods). We let *K* vary from 2—9, and for each *K*, computed the Adjusted Mutual Information (AMI) of the clustering with the CMS labels. k-means clustering only reproduces the CMS subtype to a significant degree for *K* values of 4—6, and only using maui (Figure 1C). This clustering analysis shows that maui factors are superior at predicting CMS labels, in a fair comparison, as k-means clustering is based on distances, whose computation does not benefit from higher dimensionality — in fact, the opposite is true^11^.

Latent factors inferred by maui are predictive, using k-means, of the CMS subtype, especially using *K*’s 4—6 (Figure 1C). In order to pick the best clustering result to focus on, we computed the log-rank statistic for significance of differential survival rates between clusters (See Methods). *K* = 6 results in the most statistically significant survival difference (*P <* 0.001, Figure 1D). Note that the CMS subtypes on their own are not indicative of survival rates in the TCGA data (*P* = 0.77), and that *K* = 4 (*P <* 0.045) and *K* = 5 (*P <* 0.019), also produce clusters with significant differential survival (Figure S2). Notably, *K* = 6 is preferable to *K* = 4 and *K* = 5, as it is able to tease out a cluster with particularly poor prognosis (cluster 3), which consists mostly of a subset of tumors with the CMS2 (Canonical) designation (Figure S2).

We also compared the ability of maui, MOFA, and iCluster+ to predict patient survival, irrespective of any clustering. We first selected, for each model, a subset of latent factors which are individually predictive of patient survival, and call those *clinically relevant* latent factors (See Methods). Using those clinically relevant latent factors, we fit a Cox Proportional Hazards regression, and computed Harrell’s c-Index^20^ (See Methods). The c-Index is a measure of prediction accuracy for censored data, with a score of 0.5 being expected by random guessing, and a score of 1.0 being perfect. maui (c=0.72) outperforms both MOFA (c=0.68) and iCluster+ (c=0.64) in this benchmark (Figure 1E).

The CMS subtyping scheme, as well as much of the work in the field, is based solely on gene expression profiles. In order to examine whether maui gives better predictions of patient survival with the addition of mutations and copy number data, we also trained a maui model based on gene expression alone. The maui model based on expression alone (c=0.69) achieves a lower score than a maui model with multi-omics data (c=0.72), even when the former is given more genes as input features (Figure 1F), indicating that data other than transcriptomes do contribute to overall maui performance. One of the advantages of maui over other methods such as iCluster+ and MOFA is that it is able to learn orders of magnitude more latent factors, at a fraction of the computation time (Table 2). In order to demonstrate the advantage of being able to fit larger models, we also trained a maui model based on 6,000 multi-omics features (see Methods), and that model (c=0.75) outperforms the smaller model (Figure 1F), demonstrating the clinical utility of learning from more input genes.

**Table 2.** Summary of methods.

| Method | Notes | # Factors | Runtime |
| --- | --- | --- | --- |
| iCluster+ | Bayesian, MCMC | 10 | ~11hrs |
| MOFA | Bayesian, variational | 20 | 20mins |
| *maui* | *Multilevel Bayesian, Stochastic Gradient Descent* | *100* | *3 mins* |

Finally, we investigated the applicability of using prior information from protein interaction networks for colorectal cancer subtyping. We and others previously incorporated gene-gene interactions using a method called network-smoothing^21,22^. Network-smoothing is done by allowing binary mutation values to diffuse over a gene network, a process which assigns non-zero “mutation scores” rather than binary mutation values, to genes which either have mutations, or interact with mutated genes. We used a gene network defining interactions between genes from the STRING-db^23^ database of protein-protein interactions. We applied the *netSmooth*^22^ algorithm (see Methods) to the mutation data prior to passing it to maui and computed Harrell’s c-Index, as above. Network smoothing mutations further improves the clinical relevance of latent factors learned when integrating multi-omics data (c=0.79), (Figure 1G).

A closer examination of the clusters reveals how closely the maui clusters resemble the CMS subtypes, and where they diverge. CMS1 is captured by cluster 2, CMS2 is split between clusters 3 and 5, CMS3 is captured by cluster 0, CMS4 overlaps with cluster 4, and cluster 1 is mixed (Figures 2A-C). A similar conclusion can be reached based on a set of molecular indicators introduced in^9^: CMS1 and cluster 2 show the hypermutated (Figure S3A), CIMP (Figure S3B), and microsatellite unstable phenotypes (Figure S3C). They also have similar mutation rates among TP53, APC, KRAS and BRAF (Figure S3D), a set of commonly mutated genes in colorectal cancers.

**Figure 2.**
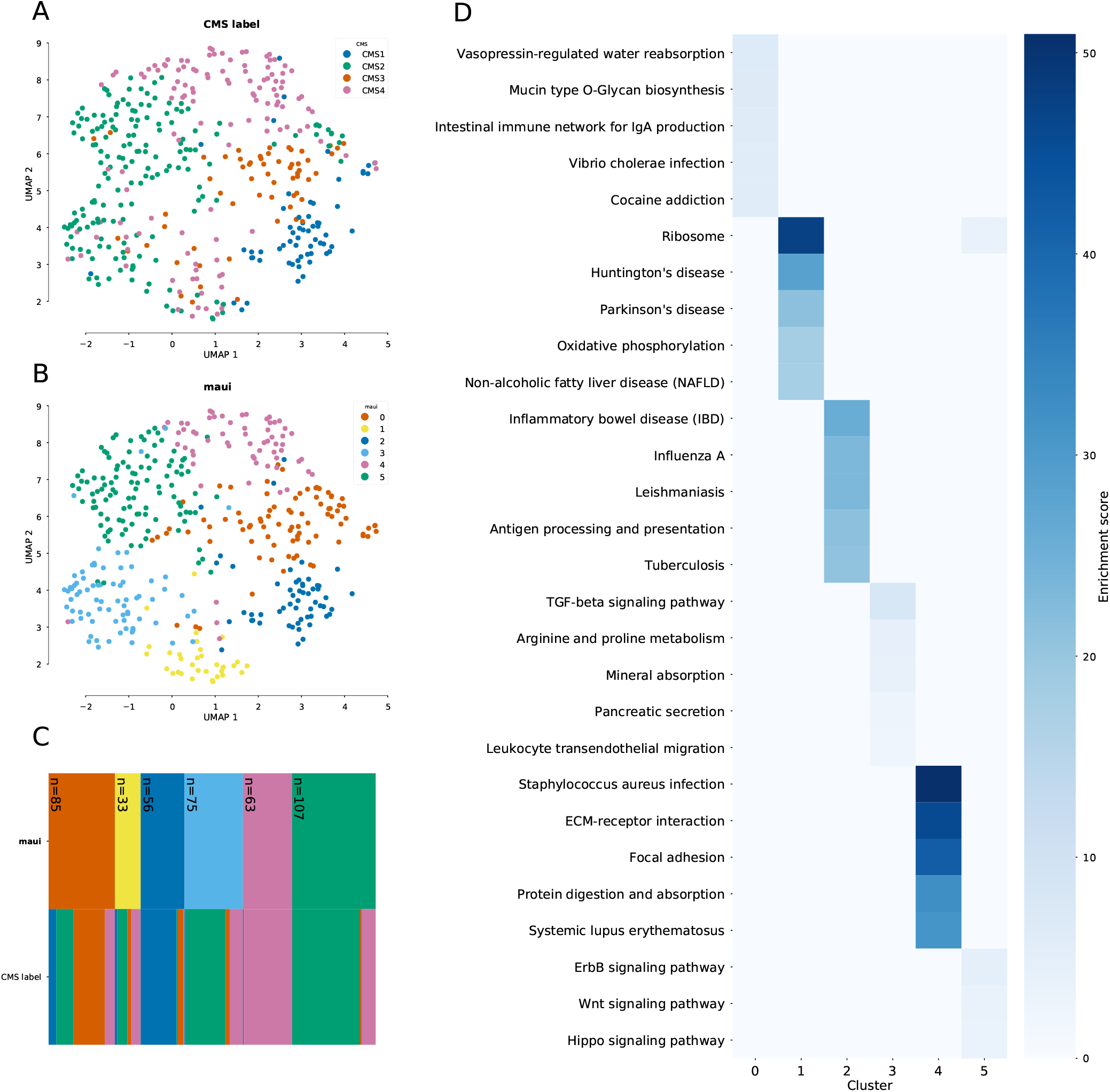
Clustering the tumors using k-means using the latent factors from maui reproduces the CMS labels closely, with the exception of CMS2 being split into two clusters, 3 and 5. **A)** UMAP embedding of tumors colored by CMS label, **B)** UMAP embedding colored by k-means clusters on maui latent factors, **C)** Cluster diagram shows the correspondence between maui clusters and the CMS subtypes: the two rows represent the different labeling schemes (maui clusters and CMS subtypes), and each column represents a sample, which is colored according with its assignment in each row. The legend in subfigures A-B applies to the color scheme in C as well. **D)** Pathways that are enriched in differentially expressed genes for each maui cluster. Clusters show a disjoint set of dysregulated pathways, underlining the different molecular phenotypes which underlie each group. Cluster 3 and 5 (which together make up the bulk of CMS2) are dominated by dysregulation of TGF-*β* signaling, and ErbB/Wnt/Hippo signaling, respectively.

Figures 2C and S3 beg the question of why CMS2 was split into two clusters (3 and 5). In order to investigate whether it is biologically plausible that the CMS group needs to be split into two, we performed a differential expression analysis, identifying marker genes for each cluster. We then ran these lists through a gene set enrichment analysis (See Methods). The top pathways associated with each maui cluster are associated with a distinct set of pathways (Figure 2D). Specifically, cluster 3 is dominated by TGF-*β* signaling and leukocyte migration, while cluster 5 is dysregulated in the ErbB, Hippo, and Wnt signaling pathways, demonstrating that these are indeed distinct groups with different molecular phenotypes. Further demonstrating this, cluster 3 presents a worse prognosis than cluster 5 (log-rank *P <* 0.001, Figure S4). Another cluster, cluster 4 (CMS4) is enriched in pathways associated with mobility and structural differences (Figure 2D), which is consistent with CMS4 displaying more stromal infiltration^9^.

### CRC latent factors are associated with processes related to tumour progression and development

Thanks to its superior computational efficiency, maui is able to infer many latent factors from multi-omics data. This creates an opportunity to select the most interesting latent factors and treat them as potential biomarkers. In order to demonstrate this, we fit Cox Proportional Hazards models^25^, fitting one regression model for each factor, as above, selecting clinically relevant latent factors (See Methods). Figure 3B shows the 95% confidence interval of coefficients for these latent factors, showing that high values for some of these latent factors are predictive of a poor prognosis (*β >* 0), while others are predictive of more favorable outcomes (*β <* 0). In general, these latent factors can be used as biomarkers with a significant prognostic value.

**Figure 3.**
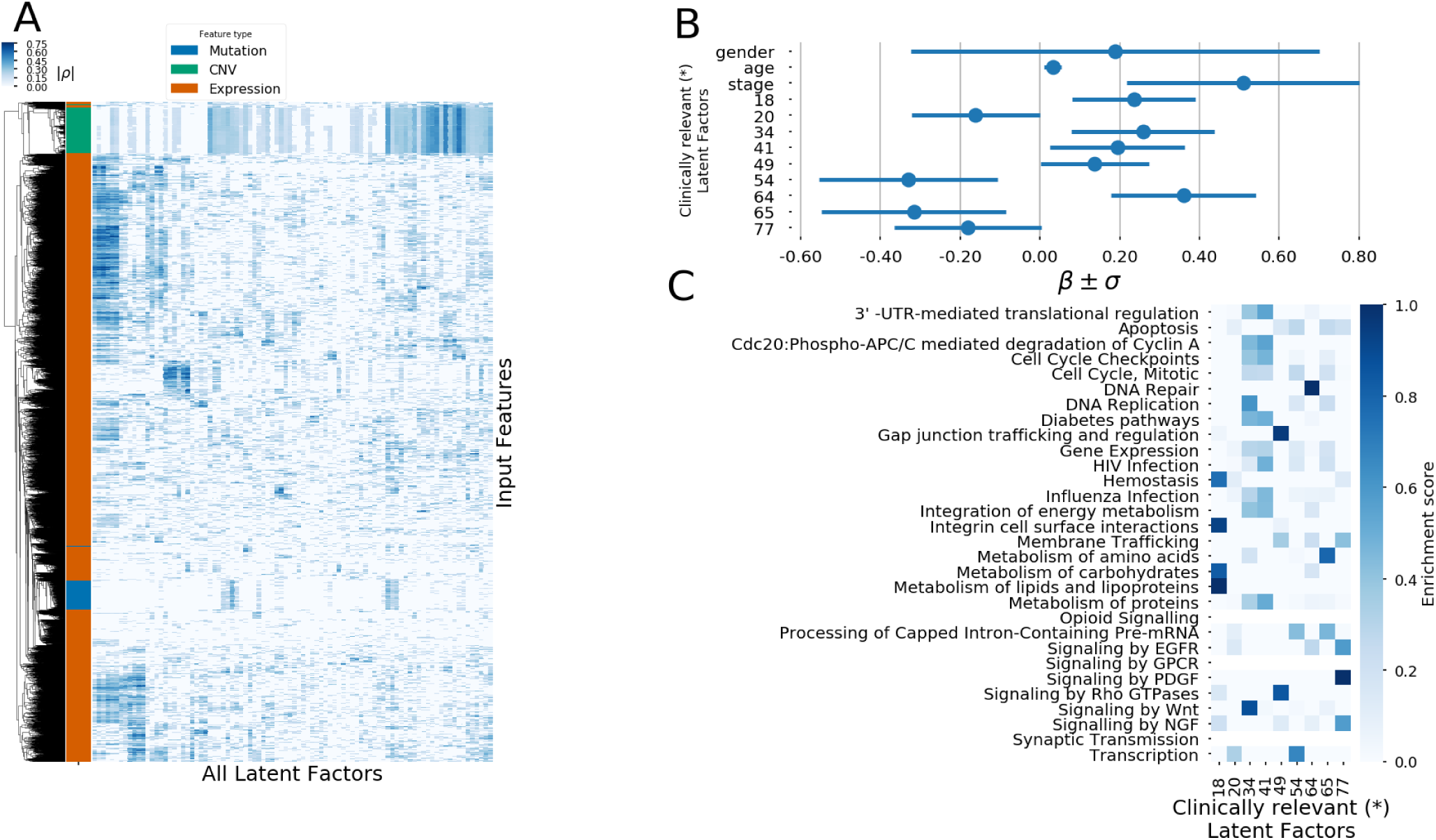
Interpretation of maui latent factors. **A)** A heatmap showing the absolute correlation coefficients of the different input genes with the latent factors. Only input features with significant correlations (*P_adj_<* 0.01, see Methods) are shown in the heatmap. The row annotation shows the type of input feature, i.e. expression value, mutation, or copy number. **B)** The coefficients in a Cox Proportional Hazards regression for factors which are clinically relevant (*) when controlling for patient age, sex, and tumor stage. Coefficients also shown for those covariates. **C)** Pathway enrichment scores for genes associated with the latent factors which carry prognostic value (have significant effects in Cox regression). (*) Clinically relevant factors are factors with a coefficient in a fitted Cox model controlling for age, sex, and tumor stage, which are statistically significantly nonzero (*P_adj_<* 0.05).

Beyond the potential to use these latent factors values as biomarkers in order to prognosticate, it is important that we be able to interpret what these biomarkers represent. maui is very powerful because it can learn highly non-linear patterns. This comes at a certain cost: the biological interpretation of the factors is less straightforward than in a linear matrix factorization approach, like PCA or MOFA. PCA and MOFA learn linear relationships between genes and latent factors, of the form *x* = *W z*, where *W* is directly available, and in it the connections between latent factors and genes. maui does not produce a straightforward, linear *W*, and so, in order to associate latent factors with input genes, we correlated input genes with latent factor values (see Methods). While most latent factors are active in the gene expression domain, most are not significantly affected by mutation data, while others capture interactions between two or more omics types (Figure 3A). By correlating latent factors with input features in this way, we can overcome the difficulties presented by the nonlinear relationships between factors and input features, and use the associations in order to find biologically relevant interpretations for neural latent factors.

When we associated clinically relevant (See Methods) latent factors with gene ids, we observed enrichment of pathways known to play a role in CRC such as Wnt signaling and other APC mediated processes (Figure 3C). In addition, one of the the most significantly survival associated factors is enriched in Neuronal growth factor (NGF) signaling associated genes. NGF signaling, which controls neurogenesis, is associated with aggressive colorectal tumours^26,27^. Survival-relevant latent factors also implicate Platelet-Derived Growth Factor (PDGF) signaling which is also associated with stromal invasion and poor prognosis for colorectal cancer patients^28,29^. Thus, in addition to using latent factors as potential biomarkers for prognosis, we can also point at the underlying biological processes that are uncovered by maui, potentially driving future drug-target studies.

### Quality assessment of CRC cell lines as models for tumors

Cancer cell lines often differ significantly in their molecular profiles from tumors, as they face a selection pressure that is different from the natural tumor micro-environment, and acquire genomic alterations necessary to adapt to cell culture^30^. As such, not all colorectal-derived cancer cell lines may be appropriate as models for tumors. Further, as cancer cell lines are maintained over time the risk of contamination and mis-labeling rises. For instance, a cell line which was originally annotated as colorectal, has been shown to be derived from melanoma^31^. Therefore, it would be beneficial if we could tell good models from ones which have diverged too much from tumors in their molecular makeup, or ones which have been mis-labeled or contaminated. To that effect, we examined 54 cancer cell lines originating in colon cancer from the Cancer Cell Line Encyclopedia (CCLE). We used maui to infer latent factor values for the cell lines, so that they could be characterized by the same latent factors as the tumors. Using latent factors, we compiled a list of nearest neighbors (See Methods) for each cell line, and counted how many of its nearest neighbors are cell lines (as opposed to tumors). The underlying hypothesis is that cell lines are ill-suited to model tumors if they are more similar to other cell lines than to tumors. We investigate this similarity in the space defined by the latent factors. About half of the colorectal cancer cell lines cell lines belong to a “cell line cluster”, meaning a majority of their neighbors are other cell lines (Figure 4A). We singled out cell lines where this proportion is above half, and found among them a mis-labeled cell line: *COLO741*, which has been shown to derive from melanoma and not colorectal cancer^ii^. This finding indicates that this method of flagging cell lines as poor models for tumors by the number of other cell lines in their neighborhood has merit.

**Figure 4.**
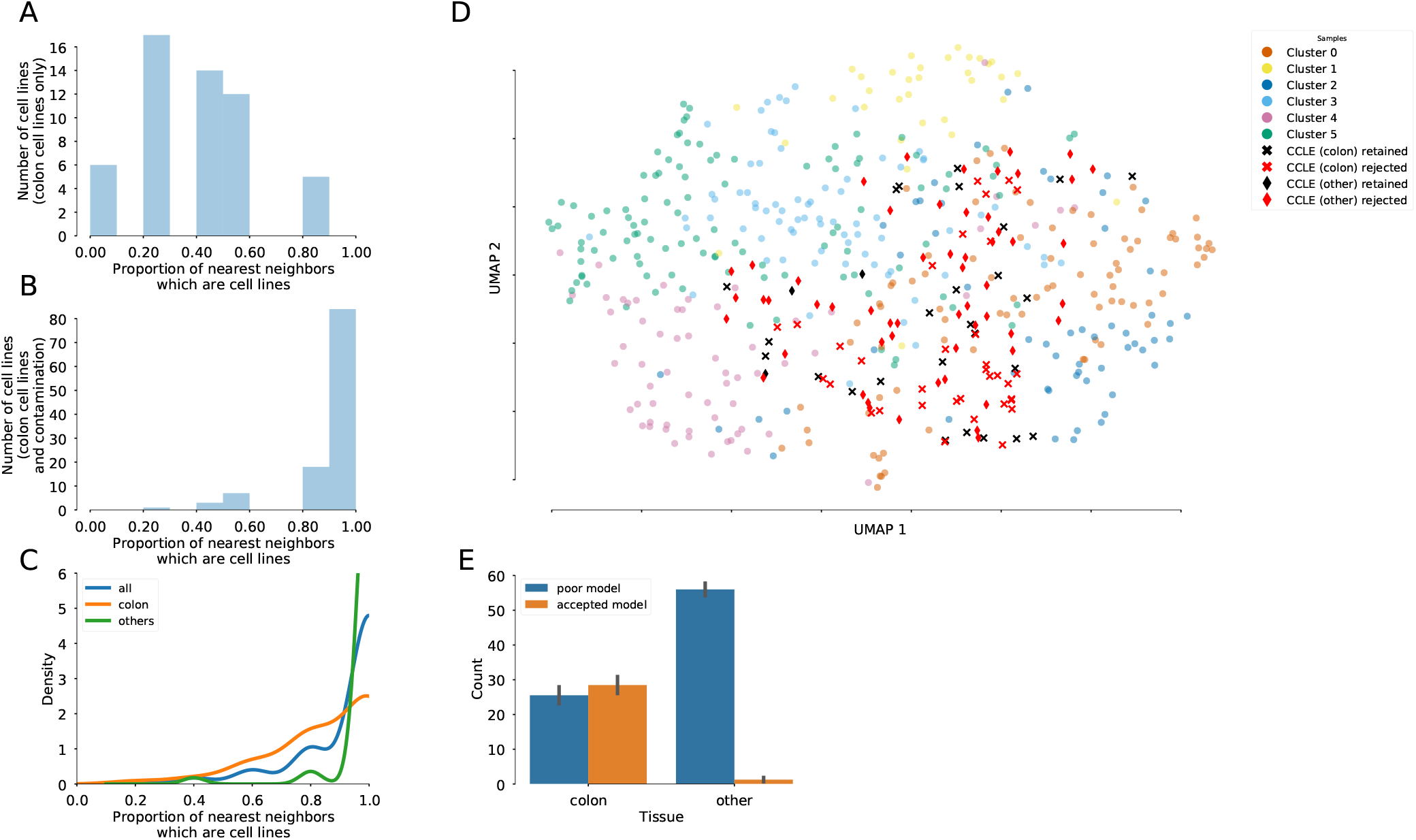
For each cell line, we compiled a list of 5 nearest neighbors in latent factor space, and counted the proportion of those nearest neighbors who are cell lines (as opposed to tumors). Cell lines whose 5 nearest neighbors are all other cell lines, are marked as bad models for tumors, as they are more similar to cell lines than to tumors. **A)** histogram of the proportion of nearest neighbors of cell lines which are also cell lines, colorectal cancers only, **B)**, histogram of the proportion of nearest neighbors of cell lines which are also cell lines, colorectal cancers and non-colorectal cell lines **C)** KDEs of the proportion of cell-line neighbors among all cell lines (colorectal and non-colorectal), broken down by tissue, **D)** UMAP embedding of tumors and cell lines. Crosses are colon-derived cell lines, diamonds are artificial contamination (non-colon derived cancer cell lines). Red cell lines are rejected, black ones are retained as good models. **E)** The proportions of colon and non-colon cell lines which are rejected because their proportion of nearest-neighbor-cell-lines is above the threshold. Nearly all non-colon cell lines are consistently rejected, as well as about half of the colon cell lines.

In lieu of knowledge of other mis-labeled or otherwise inappropriate colon-derived cancer cell lines, we artificially contaminated the data set by adding to it random sample of 60 non-colon cell lines, under the assumption that these are ill-suited to the study of colorectal cancer tumors^iii^. We then repeated the exercise of counting nearest-neighbors-that-are-cell-lines, with the additional cell lines. The introduction of these true positives^iv^ shows that more cell lines belong to a “cell line cluster”, i.e. the majority of their neighbors are other cell lines (Figure 4B), and nearly all non-colon derived cell lines have 5 of their 5 nearest neighbors be other cell lines, while the same is not true for colon-derived cell lines (Figure 4C). Hence, we designated cell lines whose 5 nearest neighbors are other cell lines, as less suitable for the study of tumors (rejected), as these appear to be more similar to other cell lines (even cell lines of other tissues) than to colorectal tumors. Cell lines that have at least one tumor among their 5 nearest neighbors, we retain as suitable models. The choice of *K* = 5 for the number of nearest neighbors is immaterial, as the method is insensitive to the choice of *K* (Figure S5). UMAP embedding of the latent factor space of tumors (with CMS labels, n=419), colorectal cancer cell lines (n=54), and non-colorectal (artificial contamination, n=60) cancer cell lines shows that most contamination cell lines are rejected, as well as some of the colon cell lines, and that the non-rejected cell lines are spread among all clusters (Figure 4D). We repeated the analysis with 100 more random draws of 60 additional contaminants, and for each such draw, rejected any cell line whose 5 nearest neighbors are cell lines. This method consistently rejects almost all known contaminants, as well as about half of the colorectal cell lines (Figure 4E). Note that it is not necessarily a mistake to reject these cell lines; being correctly labeled as originating in colon cancer does not guarantee a cell line to be a good genomic model for a tumor. Moreover, being more similar to non-colon-derived cancer cell lines than to CRC tumors, is certainly an indication that a particular cell line might not be suitable as a model for colorectal cancers, and the evidence suggests this method successfully rejects almost all known contaminants, indicating that the rejected colon cell lines are likely also poor models. Among the colorectal cell lines, CL40, SW1417, and CW2 are the best candidates, deemed most suitable as models for CRC tumors (Figure 5). On the same scale, cell line COLO320 was one of the lowest ranking cell lines. COLO320 lacks mutations in major driver genes in CRC such as BRAF, KRAS, PIK3CA and PTEN, and it has actually a neuroendocrine origin^32,33^. Therefore, COLO320 is possibly a poor model for CRC.

**Figure 5.**
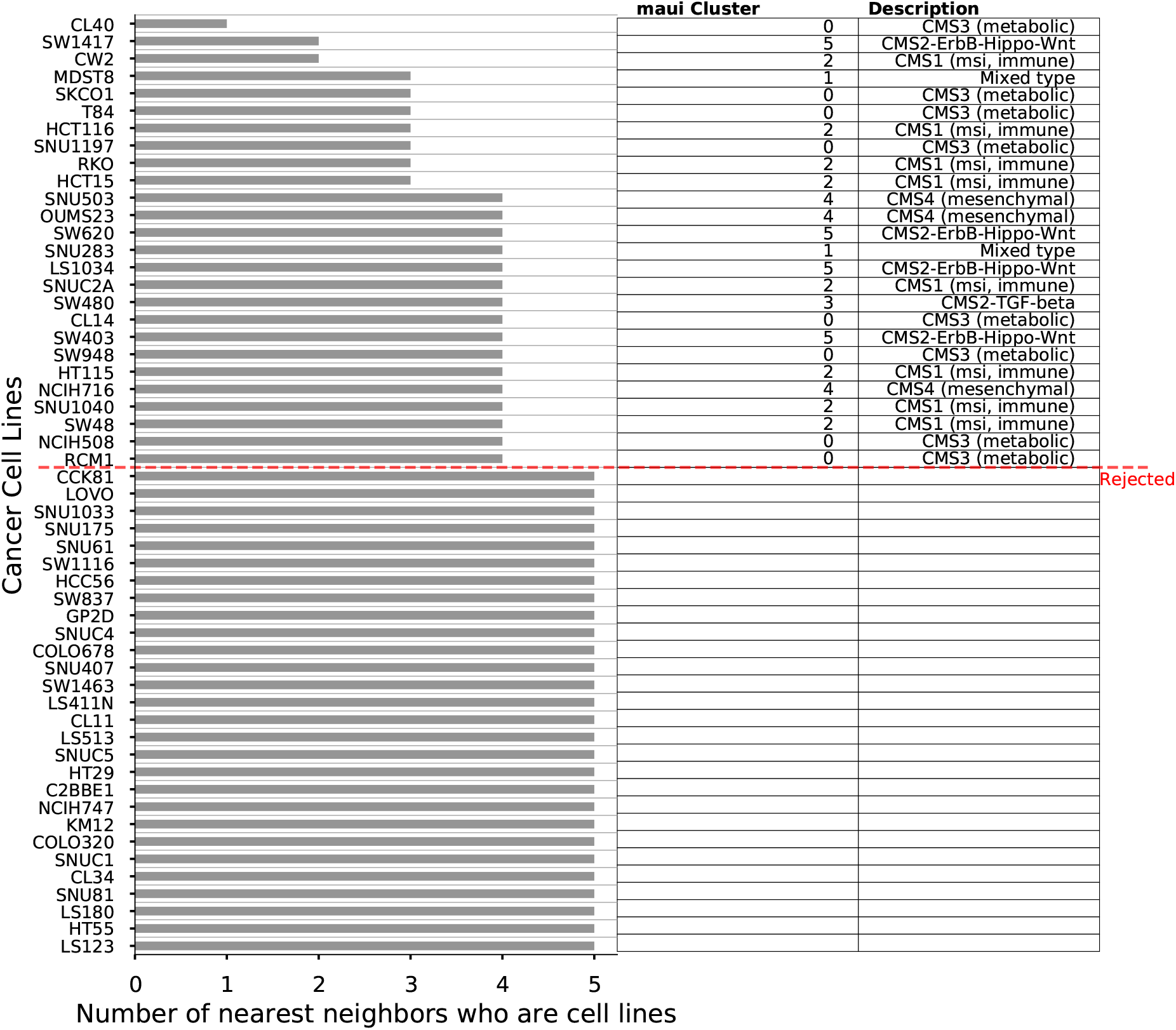
For all colon-derived cell lines, we compiled lists of their 5 nearest neighbors. The barplot shows how many of those 5 were other cell lines. Cell lines where all 5 nearest neighbors are other cell lines are rejected, those having at least one nearest neighbor who is a tumor are kept and assigned to clusters, as shown in the table on the right.

### A complete subtyping scheme for CRC and appropriate cell lines for the study of each subtype

The Consensus Molecular Subtyping (CMS) scheme^9^ is incomplete as it leaves many tumors without a CMS label. we used maui to assign subtypes to the remaining non-CMS tumors by repeating the clustering analysis, and including also tumors that don’t have a CMS designation, as well as cancer cell lines. By also including the cancer cell lines which were deemed to be suitable models (see above), we also assigned CRC subtypes to the cancer cell lines. Here, we present a novel subtyping scheme for CRC, which covers the whole TCGA cohort (including tumors without a CMS designation), as well as an association of CRC cell lines with these subtypes. The non-CMS samples are distributed roughly according to cluster size, as is to be expected for samples that lack a consensus definition (Figure 6A-B), and all clusters have at least one associated cell line (Figure 6C, Table 3). The cluster with the most associated cell lines is cluster 2 (CMS1, MSI), which consists of hyper-mutated tumors with low chromosomal instability, and indeed the cell lines that we matched with cluster 2 show the same characteristics (Figure S6), again indicating that latent factors capture patterns which are important to cancer biology. We hope that this can be a useful resource for future drug discovery studies in colorectal cancers.

**Figure 6.**
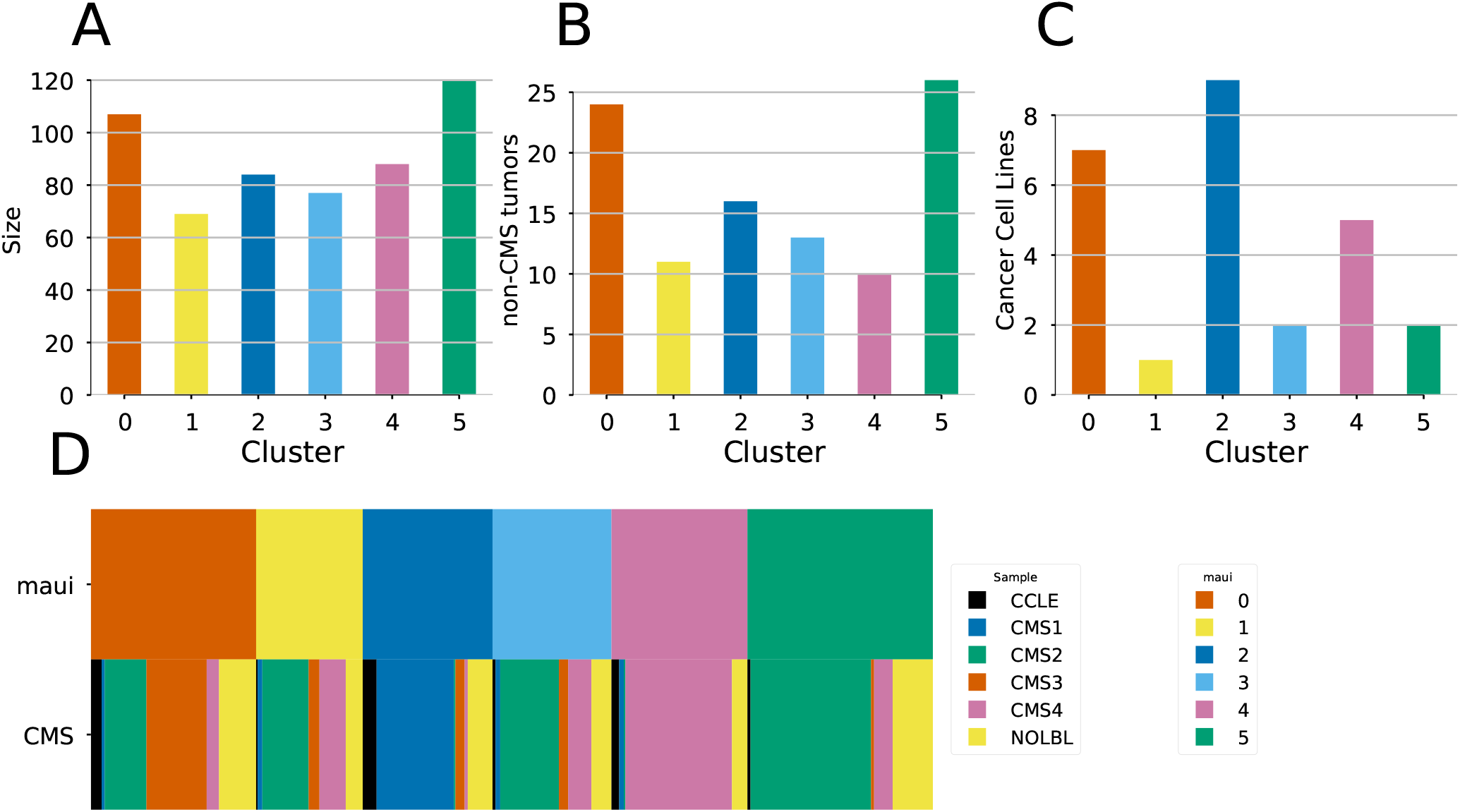
**A)** The sizes (number of samples) of the clusters, **B)** The number of non-CMS tumors assigned to each cluster, **C)** the number of cancer cell lines associated with each cluster **D)** Cluster diagram shows the correspondence between maui clusters and the CMS subtypes: the two rows represent the different labeling schemes (maui clusters and CMS subtypes), and each column represents a sample, which is colored according with its assignment in each row. The *NOLBL* samples without a defined CMS subtype are distributed among all clusters, as are cancer cell lines (CCLE).

**Table 3.** maui clusters and the cancer cell lines associated with them.

| Cluster | Description | Cell lines |
| --- | --- | --- |
| 0 | CMS3 (metabolic) | SW948, CL14, SNU1197, RCM1, NCIH508, CL40, T84, SKCO1 |
| 1 | Mixed type | SNU283, MDST8 |
| 2 | CMS1 (msi, immune) | CW2, HT115, SNU1040, HCT15, SW48, HCT116, RKO, SNUC2A |
| 3 | CMS2-TGF-beta | SW480 |
| 4 | CMS4 (mesenchymal) | OUMS23, SNU503, NCIH716 |
| 5 | CMS2-ErbB-Hippo-Wnt | SW403, LS1034, SW620, SW1417 |

## Discussion

Colorectal cancer (CRC) is a heterogeneous disease, with different subtypes being driven by different kinds of genomic alterations, e.g. hypermutated tumors, tumors showing chromosomal instability, etc. Multi-omics data analysis has the potential to increase the understanding of different subtypes of the disease, and new methods which scale computationally are necessary as the amount of available data increases. Apart from stratifying patients into clinically relevant subgroups, it is necessary to find potential drug targets specific to each subtype. Most drug target discovery studies use cancer models such as cell lines, organoids, or xenografts, and it is thus necessary to match these cancer models to the appropriate subtype in each study, or if a cancer model is inappropriate for the study of any subtype, to be able to flag it as such.

We have developed an autoencoder-based method, called **maui**, for integrating data from multi-omics experiments, and demonstrated it using RNA-seq, SNPs and CNVs. maui infers latent factors which explain the variation across the different data modalities. The latent factors inferred by maui capture important biology such as different gene expression programs, mutational profiles, copy number profiles, and their interactions. We showed that, using maui to learn latent factors in multi-omics data, we get latent factors which are predictive of previously described CRC subtypes (the Consensus Molecular Subtypes, CMS). maui outperforms the other methods we benchmarked it against, namely iCluster+ and MOFA. maui also outperforms MOFA and iCluster+ in survival prediction regardless of the CMS subtypes. From the standpoint of computational performance, maui can extract more latent factors from larger datasets, at a fraction of the computational cost of both iCluster+ and MOFA, making maui better suited to the analysis of the larger datasets we expect to see more of in the future. We have shown that maui is able to leverage its computational efficiency to learn from larger data sets, containing more genes, to produce latent factors which are more predictive of patient survival. These latent factors also produced a novel classification for CRC. While this novel classification reproduced the CMS nearly perfectly, it revealed that one of the CMS subtypes, CMS2, is in fact two distinct tumor subtypes, with different survival characteristics, and different underlying gene expression programs. These results show that an unbiased selection of more input genes, rather than restriction to known markers or driver mutations, increases prognostic value. Our results support the idea that passenger mutations as well as driver mutations could have an effect on cancer prognosis^34^.

The latent factors can also be individually associated with genes, as well as by their individual relevance to survival prediction. When we performed a pathway analysis on the latent factors which are most predictive of patient survival, we observed enrichment of pathways which are known to play a role in CRC, such as WNT signaling and other APC-mediated processes, NGF signaling, and PDGF signaling^35^. While the association of latent factors to individual genes is not as straightforward using maui as it is using matrix factorization methods, the relevance of the implicated pathways is promising. We also proposed a way to use the latent factors learned by maui to predict the fitness of cancer cell lines as models for CRC generally, as well as for specific subtypes. In order to address the first question, we hypothesized that cell lines which are poor models for the study of CRC tumors will show higher similarity to other cell lines than to CRC tumors. By including non-colorectal cancer cell lines in the sample and checking if a cell line is more similar to other cell lines than to CRC tumors, we correctly predict that 98% of non-colorectal cancer cell lines are poor models for CRC. The method also predicts that approximately 45% of the colorectal cell lines are poor models for CRC, a prediction which still needs to be validated by new experiments, although the method reliably rejected previously known inappropriate cell lines such as COLO741 and COLO320^31–^t172^33^. The rejected cell lines may still be used to study genetic interactions etc., but their utility in studies of e.g. adaptive drug response may be limited. On the other hand, SW480 and SW620 cell lines that are predicted to be a good match for CRC show similar drug response to clinical trials on KRAS mutant tumours^36^.

By including the predicted appropriate cell lines in the clustering analysis, we assigned CRC subtype-specific cell lines, a finding with far reaching potential for subtype specific drug trials. One of the clusters (cluster 2, CMS1) consists mainly of hyper-mutated tumors with low CIN, and the cell lines we matched with that cluster using maui, share those same characteristics; matching such characteristics has been a standard way to find disease-specific cell lines^37^, and this shows that maui cell line matches also preserve this desired behavior. We hope in the future it can be tested whether our approach to predicting fitness of cancer cell lines as models for tumors can be verified, and extended to other cancer models, such as organoids and xenografts. In that way, maui could become an indispensable part of drug discovery pipelines and speed up new therapeutics.

The CRC subtypes we used as a starting point for this study were defined based on gene expression profiles alone. As we wanted to use multi-omics data to refine these subtype definitions, we were limited to a subset of the tumors used in the CMS definition. We used only samples from the TCGA which had measurements for both gene expression, mutations, and copy numbers (n=519), while the CMS study used a larger cohort (n=4,151) and only gene expression profiles. Consequently, it is unclear whether the splitting of the CMS2 subtype into two clusters which we have proposed above would hold when presented with a larger dataset. Only once a larger multi-omics dataset is available will this question be answered.

While the autoencoder architecture of maui is able to do inference in larger data at a fraction of the time compared with matrix factorization methods such as MOFA and iCluster+, the resulting model is more challenging to interpret biologically, i.e. linking genes with latent factors is not as straightforward as in matrix factorization. We have proposed to solve this by using correlations of the input genes and the latent factors, picking the most significant ones heuristically. While we were able to show that such latent factor—gene relationships capture meaningful cancer biology and recapitulate known associations between dysregulation of certain pathways and patient survival, this method is potentially less robust than matrix factorization to these associations, and might require more user involvement in the analysis pipeline.

In this study we have developed a deep learning based multi-omics integration method (maui) and shown that it can be used to define clinically relevant subtypes of CRC, as well as predict the fitness of cancer cell lines as models for the study of tumors, and an association of cell lines to CRC subtypes. The latent factors inferred by maui are also interpretable in biological context, and predictive of patient survival, which enables the associations between underlying oncogenic processes, and patient survival. We benchmarked maui against two state-of-the-art methods for multi-omics data integration, and showed that not only is it more effective in defining clinically meaningful subtypes, it also does so with superior computational efficiency. Being orders of magnitude faster will enable maui to be used in studies with larger cohorts and more omics types, as these experiments become more abundant in the future. Further, maui is a general tool for multi-omics integrations, and may be used outside of the cancer context as well, in basic biology studies employing multiple genomic assays.

## Materials and Methods

### Data

We obtained data for tumors from the TCGA-COAD (n=389) and TCGA-READ (n=130) project designations of the Genomic Data Commons^v^ using the *TCGAbiolinks* R package^38^. We downloaded the CMS annotations for the TCGA tumors from the Colorectal Cancer Subtyping Consortium (CRCSC)^vi^. Table 1 summarizes the subtype information. The gene expression data (mRNA) is HTSeq - FPKM. Mutations were downloaded as MAF files, filtered to include non-synonymous mutations only, and represented as a binary mutation matrix where *m_i_j* = 1 if and only if gene *i* carries a non-synonymous mutation in sample *j*.Copy number alterations are GISTIC calls by gene, represented as a real-valued matrix where *c_i_j* is the GISTIC segment mean for the segment containing gene *i* in sample *j*.

Cancer Cell Line Encyclopedia data was obtained from the CCLE portal^vii^, and is the same data types as the TCGA data, with the exception that transcriptome profiles are RPKM-normalized and not FPKM. We considered 54 cancer cell lines originating from the colon.

We considered only tumors (from TCGA) and cancer cell lines (CCLE) which have “complete data”, that is, available measurements in all three assays: gene expression, SNVs, and CNVs.

We used gene-wise MAD statistic, computed directly on the raw data described above, in order to select the most informative genes. For the comparisons with MOFA and iCluster+, we used the 1,000 genes with the highest MAD for gene expression, 200 for mutations, and 100 for copy number alterations, for a total of 1,300 input features. We selected the features so strongly in order to make a comparison against iCluster+ viable, and with a larger feature space the runtime would become untenable (Table 2).

For the final clustering analysis, we used a larger feature space, with 5,000 gene expression values, 500 mutations and 500 CNVs for a total of 6,000 features, taking advantage of maui’s neural network architecture which allows for larger feature spaces to undergo feature selection as part of the training.

We fit the autoencoder using all TCGA samples, both with and without a CMS label (n=519, Table 1) as well as colon-derived cancer cell lines (n=54), for a total training set size of 573. For the analysis that depends on a CMS label being available, the input features were the latent factors, and the samples only those TCGA samples with a well-defined CMS label (n=419, See Table 1).

All input features were scaled and centered prior to feeding to the neural network, using batch normalization. Prior to this scaling, mutation data was binary, CNV data GISTIC calls, and gene expression counts were RPKM/FPKM values which were scaled and centered. TCGA and CCLE gene expression matrices were first scaled and centered individually, and then concatenated and scaled jointly, in order to filter out the “batch effect” of CCLE vs. TCGA data and enable mapping of tumors and cell lines to the same space. This means that, for a trained maui model, when new, unseen samples are to be mapped onto the latent factor space, they must first be normalized in this way to fit the distribution of the training data.

### Network-smoothing of multi-omics data

We applied *netSmooth*^22^ to the binary mutation matrix prior to feeding it into the neural network of maui. The method uses the protein-protein interactome (PPI) in order to *smooth* noisy molecular assays, in effect incorporating prior data from countless previous experiments, in order to improve the signal-to-noise-ratio. The intuition behind the method is that genes seldom act alone, and genes in close neighborhoods in the PPI are expected to behave similarly. For instance, interacting proteins tend to be co-expressed^39^, and somatic mutations or amplifications/deleteions in interacting may lead to similar dysfunctions.

The algorithm is a simple *Random Walks with Restarts* diffusion process on the PPI, described by the iterative process

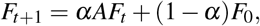

where *F* is a data matrix (gene expression, mutations, etc.), *A* is the degree-normalized adjacency matrix of the PPI, and (1 *− α*) is the restart rate. The process is guaranteed to converge, and has a closed-form solution

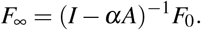

In order to pick the optimal *α* value, we performed a grid search over a range between 0 and 1, and picked the lowest *α* value within 1 standard deviation of the highest score on the Harrel’s c-Index benchmark (Figure 1E-G).

### Latent factor model for multi-omics data

Starting from different data matrices *x_i_* from different modalities, we call the full multi-omics data set *x* = [*x*_1_,*x*_2_,…,*x_m_*].

We define a generative model *x ~ p*(*x|z*). Graphically, our model looks like Figure 7a, a Bayesian latent variable model where the variation in the data *x* is explained by the variation in a smaller set of latent factors, *z*. In order to infer the latent variables *z ~ p*(*z|x*), as *p*(*z|x*) is generally intractable, we proceed with a variational Bayes framework, i.e. approximating *p*(*z|x*) *≈ q_θ_* (*z|x*), where *q_θ_* (*z|x*) is a simple class of distribution, and minimizing the Kullback-Leibler divergence *D_KL_*(*q_θ_* (*z|x*)∥*p*(*z|x*)). This is equivalent to maximizing the Evidence Lower Bound (ELBO)^40^:

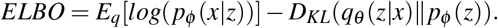

**Figure 7.**
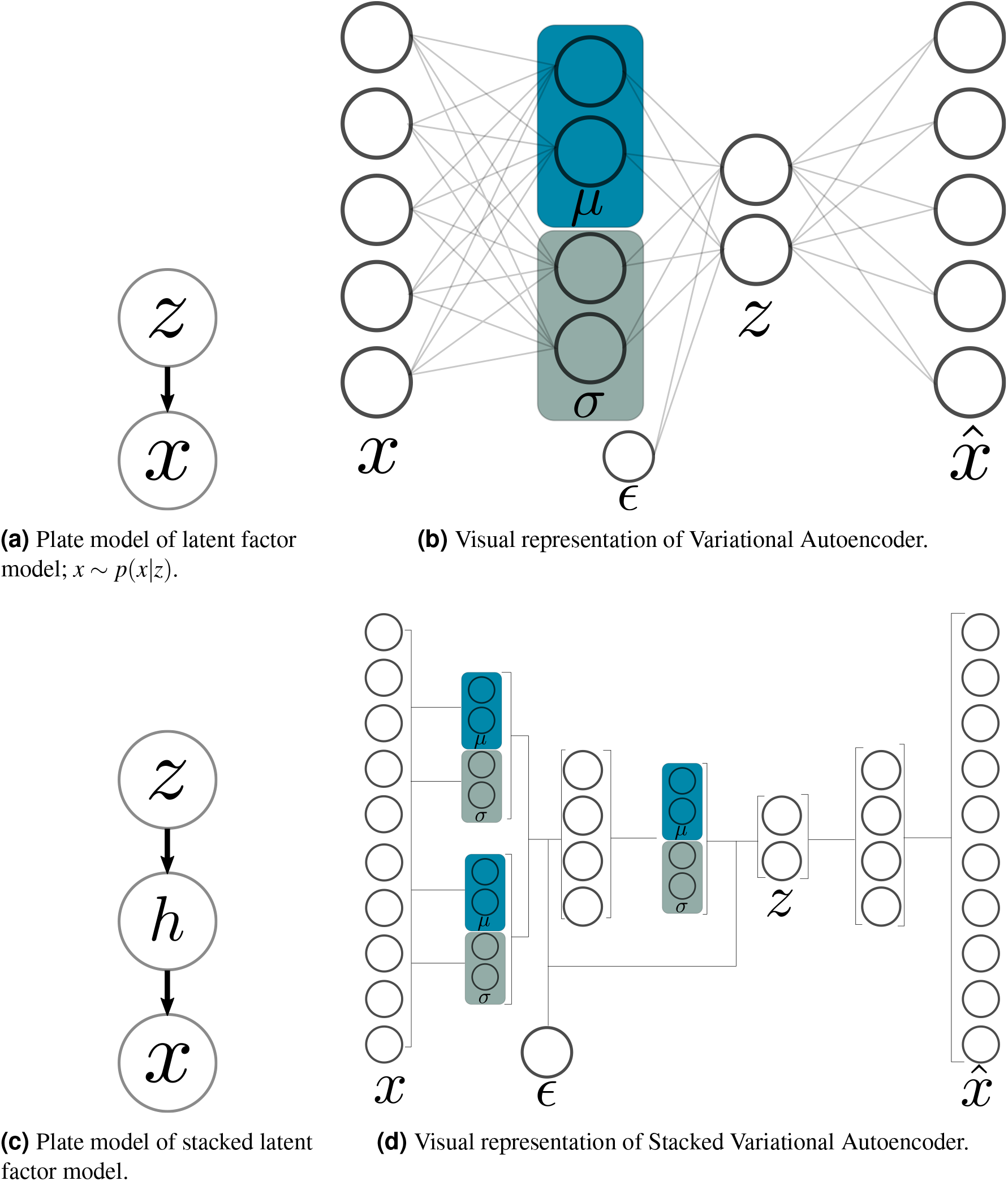
Graphical models and neural network schematics of corresponding Autoencoders. a, b: latent variable model. c, d: multilevel latent variable model.

We follow^41^ and re-parameterize 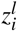 as

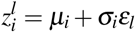

where

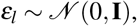

which allows us to construct the Autoencoder shown in Figure 7b.

The first half of the autoencoder, leading from *x* to *z* (the “encoder”) is a neural network which will be trained to compute *q_θ_* (*z|x*), that is, *θ* denotes the weights of the encoder network. The second half, the “decoder” network, is a neural network which will be trained to compute *p_ϕ_* (*x|z*), so *ϕ* denotes the weights of the decoder network. Thanks to the reparametrization of *z*, the path from *x* to *x*̂ is differentiable, via backpropagation, in *θ* and *ϕ*, and thus we can use gradient descent to optimize a loss function that is differentiable in *θ* and *ϕ*.

Setting the loss function of the neural network to the negative ELBO

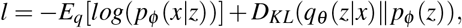

we see that the first term is equivalent to the cross-entropy reconstruction loss, and the second term, the KL-divergence between *q_θ_* (*z|x*) and the prior *p_ϕ_* (*z*) can be seen as a regularization term, which will push the *z*’s to their prior distribution.

### Stacking autoencoders

The Variational Autoencoder described above is for a one-layer bayesian framework, i.e. Figure 7a. But Autoencoders may be stacked^42^ to produce deeper neural network architectures. Deep architectures have more than one layer of nonlinearities, and can thus more compactly capture highly nonlinear functions. We introduce a hidden layer to our Bayesian latent variable model (Figure 7c).

Using the reparametrization trick as above, and specifying the full loss function, inference in the generative model (Figure 7c can be done by backpropagation in the stacked variational autoencoder model (Figure 7d).

### Model regularization

Deep neural networks have many parameters, making them very flexible. This flexibility, however, comes at a cost—deep models are prone to over-fitting: the generation of models which explain the training data well, but generalize poorly to new data. In addition, deep nets are prone to producing complex relationships between many variables. In the case of a latent variable model, that means latent factors that change with the variation of any of a large number of input features, a property which makes the task of interpreting the biological meaning of those latent factors difficult. In technical terms, we wish to enforce sparsity in *q_θ_* (*z|x*), so that each latent factor will depend on fewer of the inputs.

In order to address the first issue of potential over-fitting, we use Batch Normalization^43^. When fitting the model, we segment the data into mini-batches, at each iteration computing derivatives and making updates to the model based on that sample. Using Batch Normalization, each feature is scaled and centered in each mini-batch. We feed all of the training examples to the model fitting procedure until the entire training set is exhausted, and then we segment it into new minibatches and repeat the process, for a specified number of epochs. This way, each time a training sample is passed to the model, it will be slightly different, which is roughly equivalent to adding noise, which has been shown to work as a regularizer in Denoising Autoencoders^44^ and prevent over-fitting. Further, Batch Normalization addresses another issue - that of Internal Covariate Shift. Internal Covariate Shift happens when the distributions of activations of internal nodes in the neural network changes while training. Reducing Internal Covariate Shift enables us to pick higher learning rates, and thus speeds up inference considerably.

The second mode of regularization, encouraging disentangled representations where latent factors depend only on a few input features, is partially achieved by the KL term in the loss function, as that penalizes distributions of *z*’s which are far from the Gaussian prior. However, it has been shown that the reconstruction loss (the first term in the loss function) is generally much greater than the KL loss^15^. We therefore use their proposed method and add a multiplier to the loss function, allowing us to weigh the relative importance of the terms:

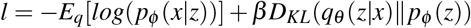

In order to ensure the network finds a good representation before it starts regularizing, we use the method proposed by^45^, where *β* is initially 0, and is gradually increased by *β* = *β* + ***κ*** until its value reaches 1.

### Model selection

The stacked VAE presented above is a class of models which are parameterized by the number of hidden units (the dimensionality of *h*), *N_hidden_*, and the number of latent factors, *N_latent_*. In order to pick the optimal model, we searched the space spanend by (*N_hidden_, N_latent_*) and computed a compound benchmark score at each point. The compound benchmark score is the average of the scores of: the AUC in the supervised CMS prediction task, the AMI in the unsupervised CMS subtype prediction task, the *−log*_10_ *p* of the multivariate log-rank test for differential survival statistics, and the c-index^20^ from the Cox proportional hazards model. The results are presented in Figure S7, where the optimal benchmark score is achieved at *N_hidden_* = 1500 and *N_latent_* = 80. Note that the general trend of the score profile is towards better scores at higher *N_latent_*, but the optimum we found breaks with that trend, indicating that this level of model complexity achieves a compromise between the most latent factors, and “the curse of dimensionality” which applies to larger latent spaces, and hurts some of the metrics.

### Model implementation

We implemented the model using Keras (v2.1.5) using a Tensorflow (v1.6.0) backend. We used Rectified Linear Units for all activations except for the last layer which is Sigmoids, for all features. We trained our network for 600 epochs using mini-batches of size 100 and *κ* = .01. We used the Adam optimizer^46^.

### Predicting CMS from latent factors using SVM

In order to quantify the correspondance between latent factors learned by different methods, adn the CMS label, we used Support Vector Machines^47^ (SVM), a supervised learning algorithm. We used a regularized SVM, picking the optimal regularization parameter using 10-fold cross-validation (CV). Then, we predict the CMS label of each sample out-of-sample, when it is in the test set, using 10-fold CV. ROC curves were computed for each class by modeling a binary outcome for each CMS label (one-vs-all). Mean ROC curves were computed by averaging the ROC of all CMS labels at each point.

### Unsupervised prediction of CMS from latent factors using k-means clustering

We benchmarked maui against MOFA and iCluster+ in the power of latent factors to predict CMS labels in an unsupervised fashion, in order to present a fair comparison between maui (70 latent factors) and MOFA (20 latent factors) and iCluster+ (10 latent factors). We used k-means clustering, as clustering based on distance metrics suffers from the “curse of dimensionality”, and does not, in general, benefit from a larger number of input dimensions (unlike supervised learning methods). To assess the ability of k-means clusters to capture the CMS labels, we ran k-means with 1,000 starts, picking the best (lowest variance) solution for each run. In addition, we applied the algorithm with *K*’s in the range of 2–9. For the cluster assignments for each *K*, we computed the Adjusted Mutual Information (AMI) of the clustering with the CMS labels. The AMI is an information-theoretic measure of the concordance between two labelings (clusterings and CMS), which accounts for chance. Higher values indicate closer relationships between labelings.

### A novel subtyping scheme for CRC with cell line associations to subtypes

The subtyping scheme presented in the results section is based on k-means clustering using maui latent factors learned from multi-omics data. We did this using a maui model trained on 6,000 input features (5,000 gene expression, 500 mutation, 500 CNV), as it is more predictive of patient survival than the one using 1,300 features (Figure 1F), and produces largely the same cluster assignments as the 1,300 gene model presented above (Figure S8).

### Association of latent factors with genomic features

The Stacked Variational Autoencoder model described above computes latent factors *z* = *f* (*x*) where *f* (*x*) is a nonlinear function which may not necessarily be well-approximated by a linear *z ≈ W x*, as in models such as MOFA or iCluster+. The architecture and depth of the neural network also makes it nontrivial to associate the input genomic features (gene expression, mutations, etc.) with the different latent factors. However, in order to make biological sense of the latent factors, it is necessary to make that association. In order to do that, we computed Pearson’s *ρ* for each latent factor with each input feature, and call a latent factor associated with an input feature if *p <* 0.001.

### Gene set enrichment

In order to identify genes associated with the different clusters, we performed a differential expression analysis using t-tests and Benjamini-Hochberg correction for multiple hypothesis testing. Genes with adjusted p value below 0.05 were called differentially expressed. In order to find out if the genes associated with latent factors (Figure 3), or with clusters (Figure 2) belong to known pathways, we used *Enrichr*^48,49^ via the python package *gseapy*^viii^. We used pathways (gene sets) defined by KEGG^50–52^.

### Survival analysis

We rely on overall survival data from the TCGA annotations for all survival analyses.

In order to assess the prognostic value of latent factors inferred by our deep learning approach, we fit a Cox Proportional Hazards model^25^,

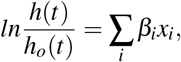

where the left hand side is the logarithm of the hazard ratio, and *x*’s are co-variates. We assess the predictive value of each latent factor separately, while controlling for the patient’s age, gender, and tumor stage at diagnosis. We compute confidence intervals for the coefficient *β* associated with the latent factor, and pick the latent factors with FDR-correction and *α* = 0.95.

In order to compare the prognostic value of different models, we compute the c-index^53–55^ and use 5-fold cross-validation^47^.

The log-rank statistics reported in Figure 1D and Figure S2 are multivariate log-rank test, under a null hypothesis that all groups have the same survival function, with an alternative hypothesis that *at least one group* has a different survival function.

All survival analysis was done using the python package *lifelines*^ix^.

### Comparing models’ survival-predictive value

In order to compare maui to MOFA and iCluster+ (as well as to a gene expression only-based maui model), we used Harrell’s C^20^ in a Cox Proportional Hazards^25^ regression model. The c-Index was computed for Cox models based only on clinically relevant factors, which we select using individual, unregularized Cox models, one per factor, while controlling for patient age, sex, and tumor stage. In those individual factor models, we used Efron’s method to compmute confidence intervals, and only kep the latent factors with statistically significant (adjusted P-value *<* 0.05) nonzero coefficients in the individual Cox models. Having selected clinically relevant latent factors from each model (maui, MOFA, iCluster+, maui-expression, maui-netsmooth), we fit a full Cox regression using those, and ran a cross validated out-of-sample c-Index calculation using regularized Cox PH regression, searching for the optimal result among the regularizers 1,10,100,1000,10000. The results reported in Figure 1F are the best regularized model for each of the methods.

### Quality assessment of CRC cell lines for modeling tumors

In order to assess the fitness of different cancer cell lines as models for tumors, we computed the pairwise euclidean distance between each of the samples (TCGA and CCLE), in the space of the latent factors derived from maui. Then, we computed, for each cell line, the proportion of its 5 nearest neighbors which are also cell lines, the working hypothesis being that cell lines that form “cell line clusters” are more cell-line like than tumor like, and likely less fit as models for tumors. We repeated the exercise considering other numbers of nearest neighbors from 1—20, at each *K* computing the true positive rate (recall), that is, 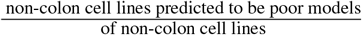, showing that the recall is near perfect for a wide range of *K*’s.

## Footnotes

i Some CRC samples do not have a consensus molecular subtype

ii In more recent versions of the CCLE annotations, this has been fixed.

iii The identities of these “known contaminant” cell lines are irrelevant, as we show later that the method works on 100 such random draws.

iv Non-colon cancer cell lines are considered true positives in the task of predicting which cell lines are poor models for CRC tumors.

v https://portal.gdc.cancer.gov

vi http://sagebionetworks.org/research-projects/colorectal-cancer-subtyping-consortium-crcsc/

vii https://portals.broadinstitute.org/ccle

viii version 0.9.4, available from PyPI https://pypi.org/project/gseapy

ix https://lifelines.readthedocs.io/en/latest/

x http://cancergenome.nih.gov/

xi https://portals.broadinstitute.org/ccle

## Acknowledgements

The work shown here is based upon data generated by the TCGA Research Network^x^ and the Cancer Cell Line Encyclopedia^xi^. We would like to thank Claudia Baldus, Gaetano Gargiulo, Vedran Franke, Bora Uyar, Wolfgang Kopp and Ella Bahry for valuable comments on the manuscript.

## Author contributions statement

AA, SH and JR conceptualized the project. AA set the objectives with input from JR and SH. The analysis presented in this manuscript, and all the software development was done by JR. SH provided additional analysis on latent factor interpretation and cell line matching. AA and JR wrote initial draft of the manuscript with input from SH. AA supervised the writing, software development and analysis. JR, AA and SH reviewed and finalized the manuscript.

## Supplementary material

**Figure S1.**
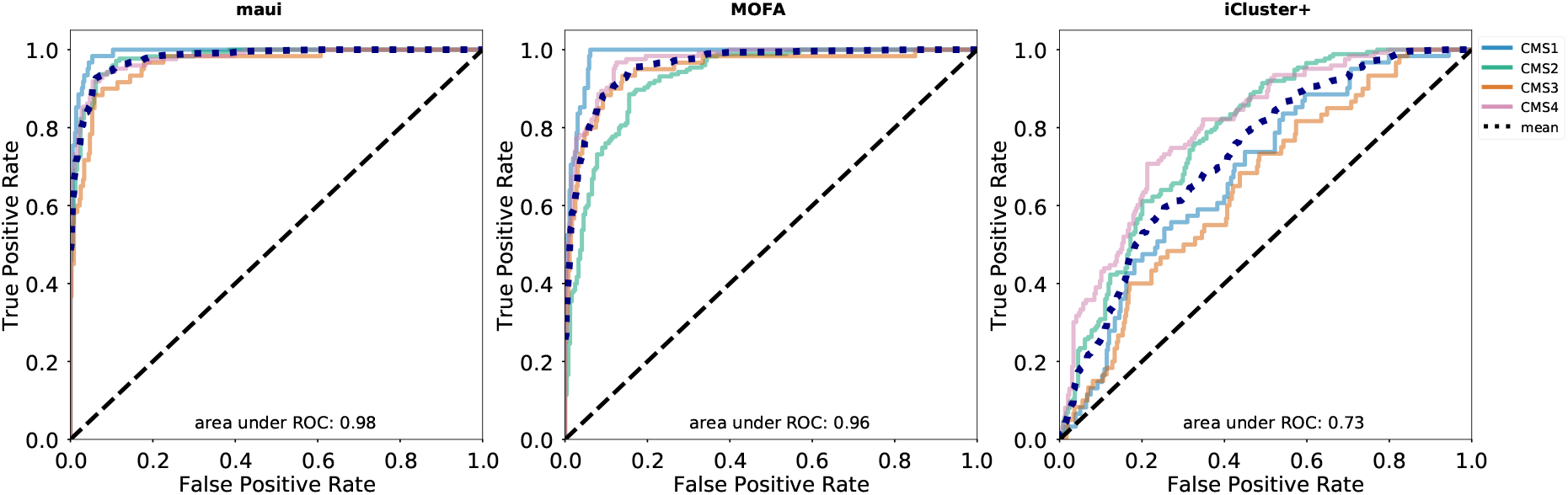
Receiver Operator Characteristic curves per class (CMS) for maui, MOFA, and iCluster+. Mean ROC curve also shown. auROC reported is the area under the mean ROC.

**Figure S2.**
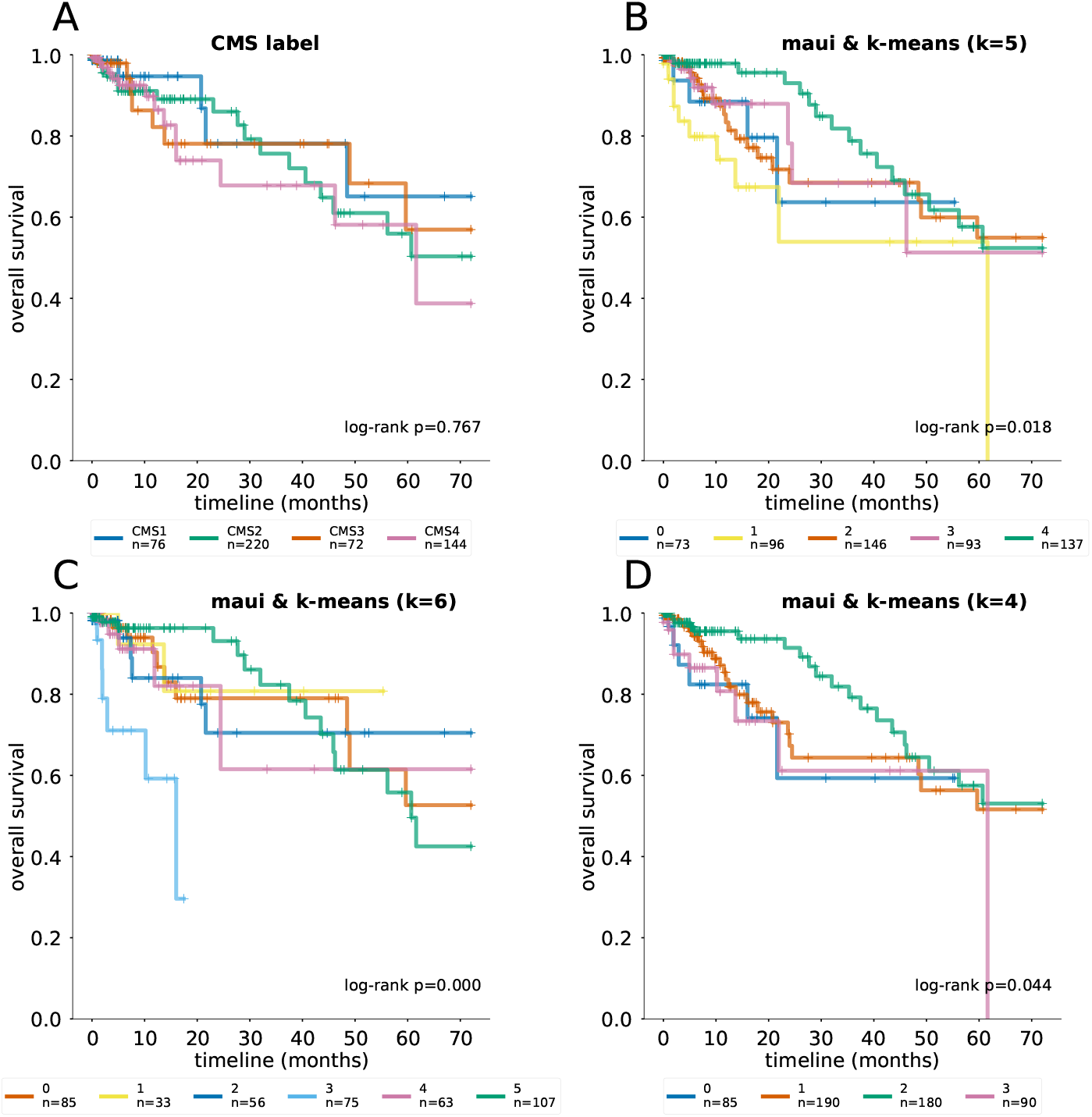
Kaplan Meier curves and log-rank tests for differential survival statistics for the CMS subtypes, as well as maui clusters using k-means with different K’s. The reported P values are from a multivariate log-rank test, under a null hypothesis that all groups have the same survival function.

**Figure S3.**
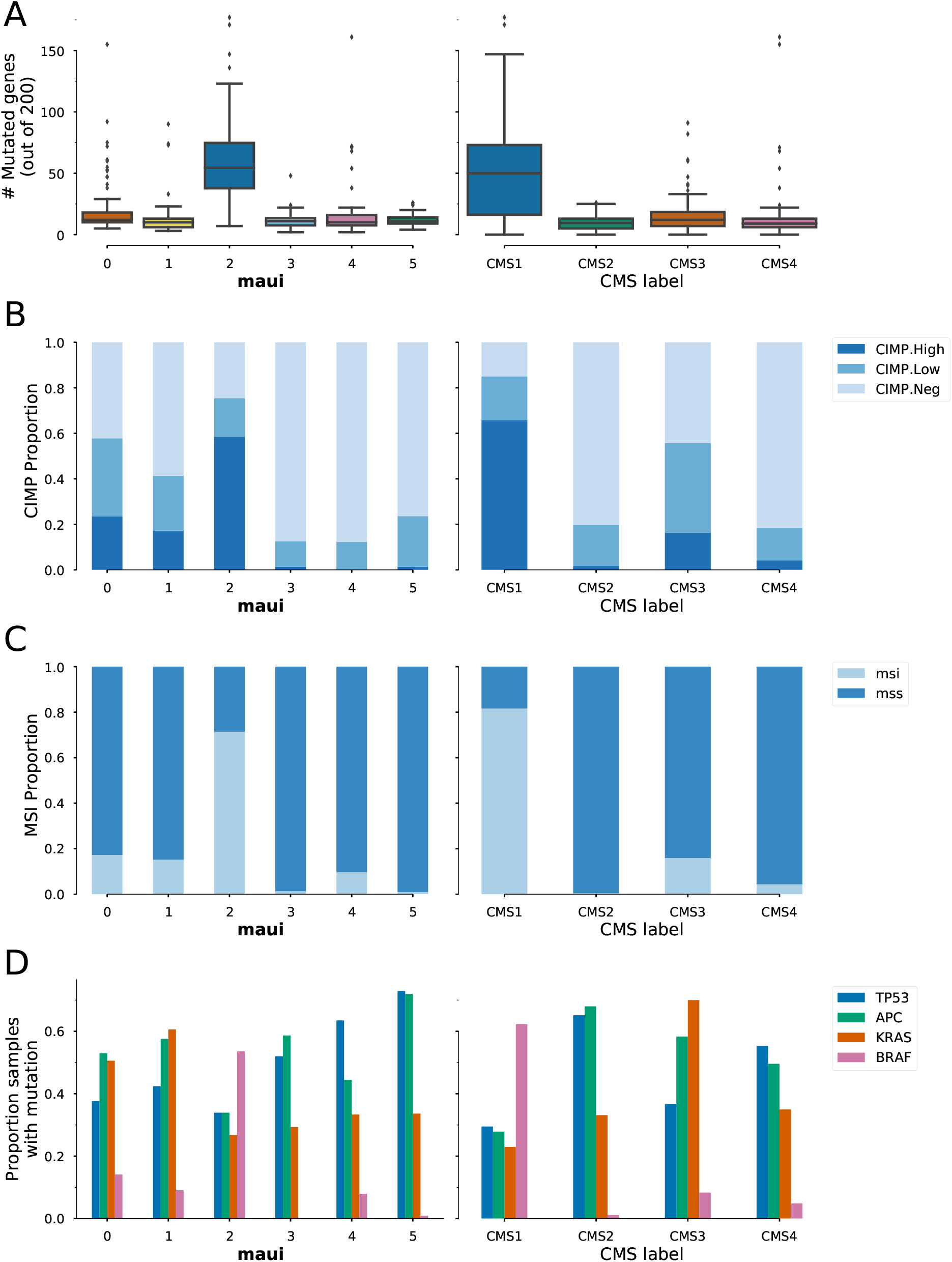
Molecular markers and their distribution in CMS subtypes (left column) and maui clusters (right column). **A)** utational load, **B)** CIMP phenotype, **C)** Microsatellite instability, and **D)** the prevalence of mutations in a key set of colorectal cancer genes.

**Figure S4.**
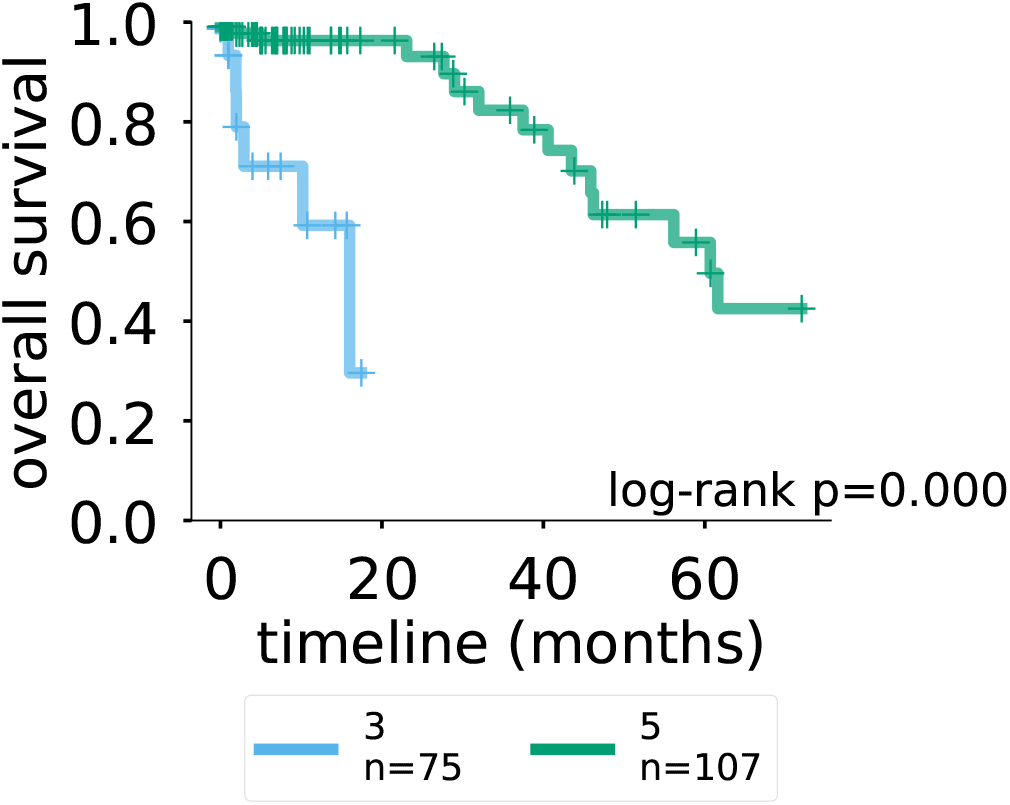
Kaplan-Meier curves for maui clusters 3 and 5. Cluster 3 appears to be more aggressive tumors with a worse prognosis (*P <* 0.001).

**Figure S5.**
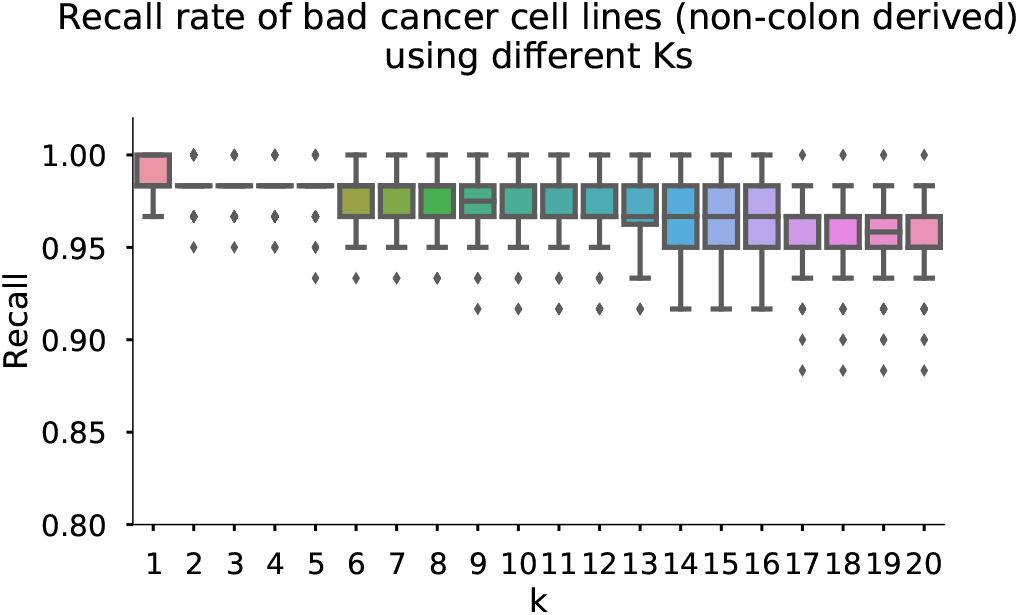
We repeated the exercise of Figure 4E, that is, adding non-colon cell lines to the mix, and calculating the proportion of each cell line’s K nearest neighbors, that are also cell lines (as opposed to tumors). Setting the threshold at 0.95, the method correctly identifies most non-colon cell lines as bad models for colorectal tumors. The recall rate is 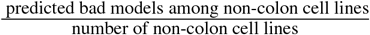, and is largely insensitive to the choice of K.

**Figure S6.**
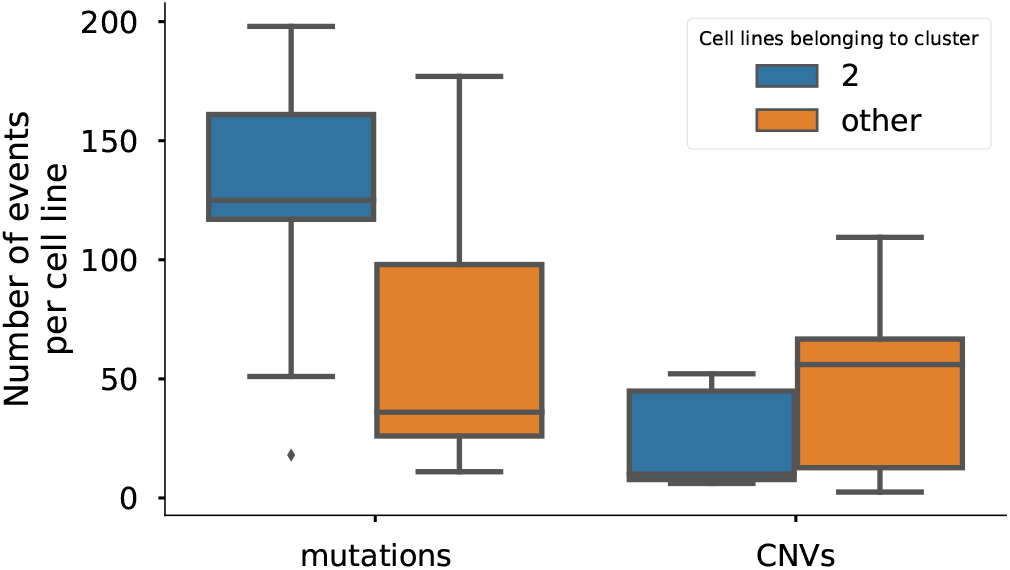
The CMS1 subtype, which is captured by maui cluster 2, consists of hyper-mutated tumors with low chromosomal instability, resulting in tumors with a large number of mutations, but low number of copy number events (Figure S3). The cell lines that we matched with cluster 2 also show the same characteristics.

**Figure S7.**
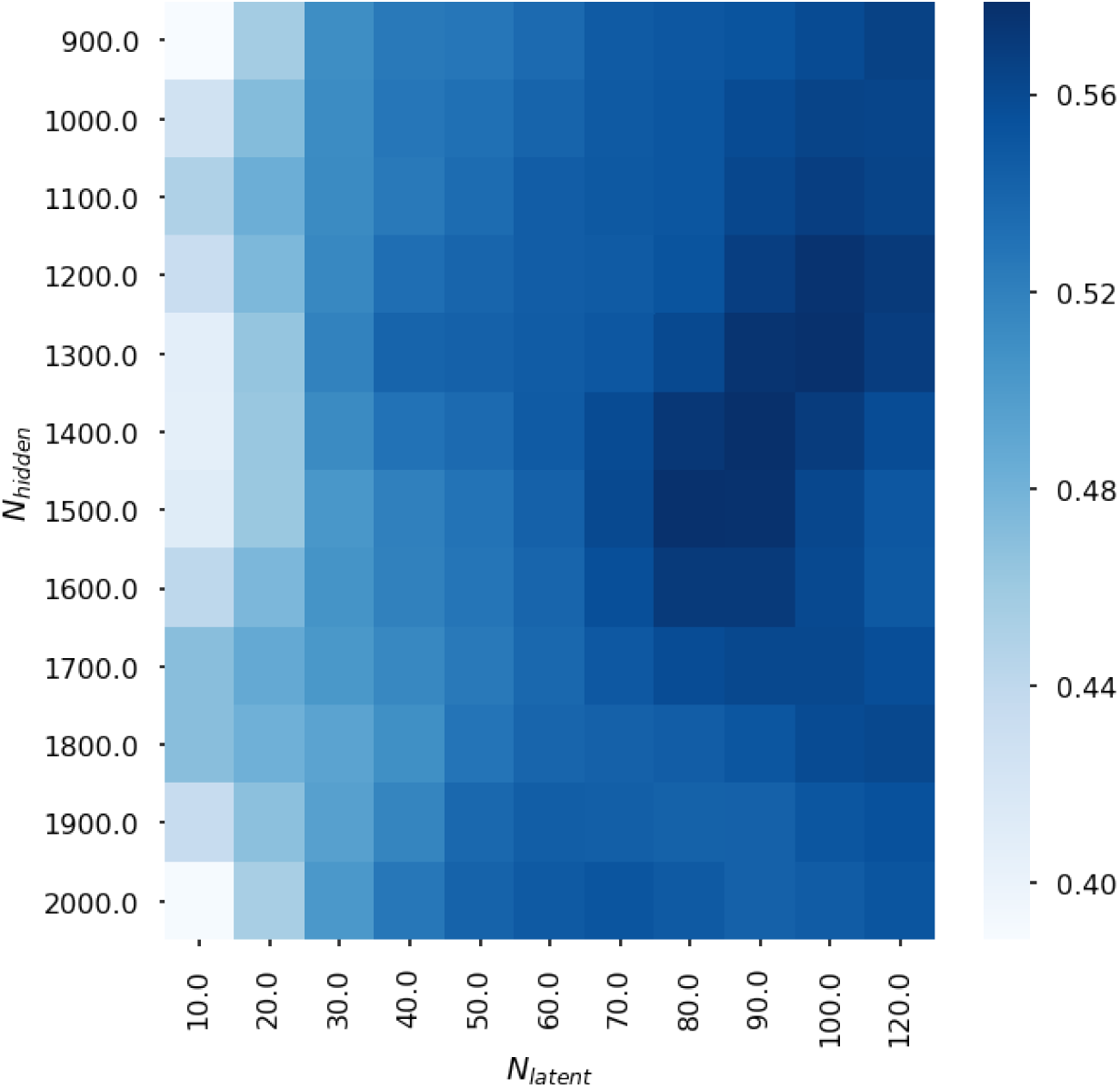
The composite benchmark score in the space defined by *N_hidden_*, the number of hidden units, and *N_latent_*, the number of latent factors in a model. The optimal parameters are *N_hidden_* = 1500 and *N_latent_* = 80

**Figure S8.**
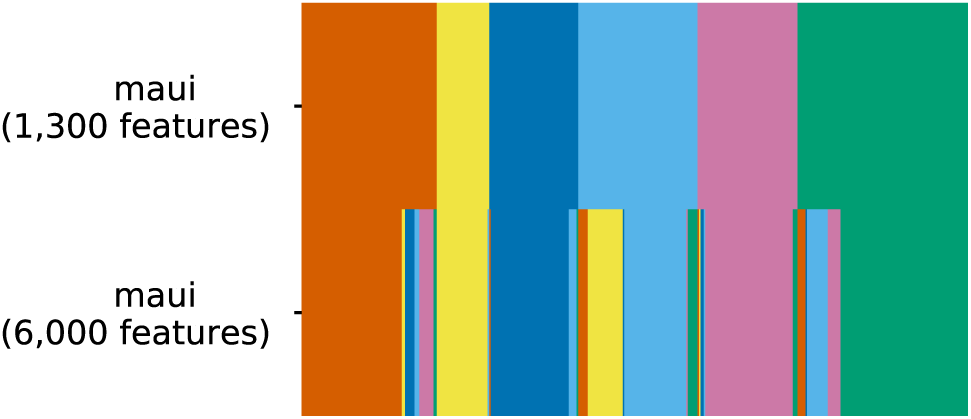
Correspondence of maui clusters when training using 1,300 genes and 6,000 genes. Each column is a sample, and they are colored by their cluster assignment. Clusters are mostly the same when using more input features, with some refinements taking place.

